# Characterization of bovine-derived H5N1 viruses expressing fluorescent and luminescent reporter proteins

**DOI:** 10.64898/2026.01.17.700109

**Authors:** Mohammed Nooruzzaman, Pablo Sebastian Britto de Oliveira, Chen Feng, Diego G. Diel

**Author notes:** Corresponding author: Diego G. Diel;. **Accession number.** All sequence data generated in this study were submitted to the NCBI BioProject (BioProject ID PRJNA1406649).

## Abstract

Highly pathogenic avian influenza (HPAI) H5N1 clade 2.3.4.4b viruses present a broad host range, with recent spillover and sustained transmission in dairy cattle reported in the United States. Replication-competent reporter viruses are critical tools that enable real-time monitoring of virus replication facilitating high-throughput screens. In this study, we engineered three recombinant H5N1 clade 2.3.4.4b reporter viruses expressing nanoluciferase (NLuc) and two fluorescent reporter proteins, miniGFP2 and UnaG within the open reading frame of the nonstructural (NS) gene of the bovine A/Cattle/Texas/063224-24-1/2024 (TX2/24) virus. All reporter viruses replicated efficiently *in vitro*, presenting replication kinetics comparable to the parental rTX2/24 virus, but exhibited smaller plaque sizes suggesting reduced cell-to-cell spread. *In vivo* infection studies in mice showed comparable pathogenicity among all four viruses, although rTX2/24-miniGFP2 and rTX2/24-UnaG exhibited decreased virus shedding relative to rTX2/24 and rTX2/24-NLuc. Virus titrations and *in situ* localization of virus replication sites demonstrated robust replication in respiratory tissues, with slightly attenuated systemic dissemination of all three reporter viruses. Fluorescent virus neutralization assays using miniGFP2 and UnaG reporter viruses accurately quantified neutralizing antibody titers in sera from naturally infected dairy cattle, consistent with wild-type virus assays. Additionally, the utility of the NLuc reporter virus for antiviral screening was validated against Oseltamivir *in vitro*. Collectively, these results establish the TX2/24-based reporter viruses as versatile and biologically relevant tools for investigating H5N1 pathogenesis to be used in serological and antiviral drug screens against H5N1 viruses.

## Introduction

Highly pathogenic avian influenza (HPAI) H5N1 viruses pose a major threat to human and animal health due to their ongoing evolution, interspecies transmission and zoonotic potential. Since their emergence in domestic geese in Guangdong, China in 1996, H5N1 viruses have undergone extensive genetic diversification, driven by their error-prone RNA polymerase and reassortment events with low pathogenic avian influenza viruses in aquatic bird reservoirs [1, 2]. Among the emerging variants, viruses belonging to clade 2.3.4.4b have become globally dominant since 2014, causing widespread outbreaks in wild birds and poultry across multiple continents [3].

In 2024, the detection of clade 2.3.4.4b genotype B3.13 H5N1 viruses in dairy cattle in the U.S. marked an unprecedented host range expansion, with confirmed spillover from infected cows to humans and other mammals-including domestic cats, raccoons, and birds-via direct and indirect contact or ingestion of contaminated raw milk [4–10]. Since the emergence of the virus in dairy cattle, at least 71 human infections have been confirmed by the U.S. Centers for Disease Control and Prevention (CDC), 41 of which involved direct contact with dairy cows, farm equipment, or unpasteurized milk [7, 11]. These events have heightened concerns regarding viral adaptation to mammalian hosts and the potential for sustained mammal-to-mammal transmission.

Understanding the pathogenesis, tissue tropism and shedding and transmission patterns of emerging H5N1 viruses is essential for accurate risk assessment and the development of effective countermeasures. Here we developed and characterized reporter H5N1 2.3.4.4b viruses expressing bioluminescent or fluorescent markers facilitate such studies [12–15]. While reporter influenza viruses have been developed for different virus subtypes, including H1N1 [13, 14, 16–18] and H5N1 [15, 19, 20], most constructs rely on avian-origin backbones or in attenuated strains, limiting their translational relevance to recently emerged H5N1 viruses.

Fluorescent and bioluminescent proteins are the two reporter systems most widely used in virological studies due to their sensitivity, stability, and ease of detection. Fluorescent proteins are ideal for observing cellular localization and *ex vivo* imaging; however, their *in vivo* signals often suffer from low intensity and interference from tissue autofluorescence [21, 22]. In contrast, luciferases are better suited for *in vivo* and non-invasive imaging due to their high sensitivity and low background signal [15, 17, 18].

In this study, we generated and characterized three recombinant reporter H5N1 viruses based on the bovine isolate A/Cattle/Texas/063224-24-1/2024 (TX2/24), representing the currently circulating clade 2.3.4.4b lineage genotype B3.13. Using reverse genetics, we inserted nanoluciferase (NLuc) and two green fluorescent proteins, miniGFP2 and UnaG, into the nonstructural (NS) segment to preserve viral replication competence [12–15]. NLuc is a 19 kDa engineered luciferase that exhibits approximately 150-fold greater specific activity (light output) than Renilla or firefly luciferases [23]. MiniGFP2 is a 13 kDa green fluorescent protein with improved photochemical stability compared to other flavin-binding proteins, enabling long-term live-cell imaging [24]. UnaG is a 15.6 kDa bilirubin-inducible fluorescent protein first discovered in 2013 in the muscle tissue of the Japanese eel (*Anguilla japonica*) [25]. We evaluated their replication kinetics and plaque morphology *in vitro*, assessed pathogenicity and tissue tropism in a mouse model, and demonstrated their application in virus neutralization and antiviral activity assays. These recombinant bovine H5N1 reporter viruses provide valuable tools for studying infection dynamics, host adaptation, and immune responses to contemporary H5N1 viruses at the animal-human interface.

## Methods

### Biosafety, biosecurity, and ethical approvals

All work involving the rescue, handling, and propagation of highly pathogenic avian influenza (HPAI) H5N1 viruses was conducted under strict biosafety and biosecurity conditions in the Animal Health Diagnostic Center (AHDC) research BSL-3 facility at the College of Veterinary Medicine, Cornell University. All animal experimental procedures were reviewed and approved by the Institutional Animal Care and Use Committee (IACUC) of Cornell University (approval number 2024-0094). Animal studies were conducted under animal biosafety level 3 facilities (ABSL-3) at the East Campus Research Facility at Cornell University. All applicable institutional and ethical regulations were followed.

### Cells

Human embryonic kidney HEK293T cells, Madin-Darby canine kidney (MDCK) cells and human lung adenocarcinoma (A549) cells were cultured in Dulbecco’s Modified Eagle Medium (DMEM) supplemented with 10% fetal bovine serum (FBS) and 1X Antibiotic-Antimycotic (ThermoFisher Scientific, Waltham, MA). Bovine uterine epithelial cells (Cal-1, developed in house at the Virology Laboratory at AHDC) were cultured in minimal essential medium (MEM, Corning Inc., Corning, NY) supplemented with 10% fetal bovine serum (FBS) and 1X Antibiotic-Antimycotic.

### Generation of recombinant HPAI reporter viruses

The recombinant rTX2/24, rTX2/24Nluc, rTX2/24-miniGFP2 and rTX2/24-UnaG reporter viruses were generated using a reverse genetics approach [26]. Briefly, full-length genome sequences of the HA, NA, M, NP, PA, PB1, and PB2 gene segments of the TX2/24 strain (H5N1 clade 2.3.4.4b, genotype B3.13; GISAID accession number EPI_ISL_19155861) were commercially synthesized (Twist Bioscience) and cloned into the dual-promoter influenza reverse genetics plasmid pHW2000 (kindly provided by Dr. Richard Webby, St. Jude Children’s Research Hospital) using *BsmBI* restriction sites (New England Biolabs). To generate the NS reporter viruses, the TX2/24 NS gene segment was modified to express proteins from a single nonoverlapping transcript. NLuc, miniGFP2 or UnaG sequence was fused to the C-terminus of NS1, and the NS1 and NEP open reading frames were separated by the porcine teschovirus-1 (PTV-1) 2A autoproteolytic cleavage site. The modified NS-NLuc, NS-miniGFP2 or NS-UnaG gene segments were commercially synthesized (Twist Bioscience) and cloned into the pHW2000 vector using *BsmBI* restriction sites.

The pHW2000 plasmids containing the seven unmodified TX2/24 gene segments together with the modified NS-reporter segment were co-transfected into a co-culture of HEK293T and Cal-1 cells using Lipofectamine 3000 (ThermoFisher Scientific, Waltham, MA). Cell culture supernatants were harvested at 96 h post-transfection and used to infect freshly seeded Cal-1 cells. Both cell lysates and culture supernatants were collected at 72-96 h post-infection to generate seed virus stocks.

Working virus stocks were prepared by inoculating 10-day-old embryonated chicken eggs via the allantoic cavity route, and infected allantoic fluids were harvested 48 h post-inoculation. Viruses from the initial rescue and two subsequent passages were sequenced to confirm the absence of unintended mutations. Viral titers were determined as 50% tissue culture infectious doses (TCID_50_) using endpoint dilution assays and calculated by the Spearman-Kärber method, and titers were expressed as TCID_50_.mL^-1^.

### Replication kinetics of recombinant HPAI H5N1 viruses

Replication kinetics of recombinant HPAI H5N1 reporter viruses (rTX2/24-NLuc, rTX2/24-miniGFP2 and rTX2/24-UnaG) were evaluated in bovine uterine epithelial cells (Cal-1), human lung adenocarcinoma A549 cells, and Madin-Darby canine kidney (MDCK) cells and compared with the parental rTX2/24 virus. Cells were seeded in 12-well plates at a density of 2.5×10^5^ cells per well and incubated at 37 °C for 24 h to reach approximately 90% confluency. Cells were then infected with one of four recombinant viruses rTX2/24, rTX2/24-NLuc, rTX2/24-miniGFP2 or rTX2/24-UnaG at a multiplicity of infection (MOI) of 0.1. Virus adsorption was carried out at 4 °C for 1 h, after which the inoculum was removed, replaced with 1 mL of complete growth medium, and plates were transferred to a 37 °C incubator. Infected cells were incubated at 37 °C, and combined cell lysate and supernatant samples were collected at 4, 8, 12, 24, 48, and 72 h post-infection (p.i.) and stored at −80 °C until analysis. The 0 h time point consisted of an aliquot of the virus inoculum collected immediately after inoculation and stored at −80 °C. Virus titers were determined in Cal-1 cells using endpoint dilution assays, with infectivity assessed by immunofluorescence assay as previously described [27, 28]. Viral titers were calculated using the Spearman-Kärber method and expressed as TCID_50_.mL^-1^.

### Luciferase reporter assay

Cell lysates from Cal-1, A549, and MDCK cells infected with rTX2/24, rTX2/24-NLuc, rTX2/24-miniGFP2, or rTX2/24-UnaG during the replication kinetics experiments were analyzed using a luciferase reporter assay to quantify NanoLuc luciferase expression. The Nano-Glo^®^ Luciferase Assay System (Promega) was used according to the manufacturer’s instructions. Briefly, 50 µL of each cell lysate was added to a white 96-well luminometer plate, followed by the addition of 50 µL of Nano-Glo assay substrate. After incubation for 3 min in the dark at room temperature, luminescence was measured using a luminometer plate reader (BioTek Synergy LX Multimode Reader).

### Plaque assays

Plaque morphology and phenotype of recombinant H5N1 viruses were assessed in bovine uterine epithelial cells (Cal-1), human lung adenocarcinoma (A549) and Madin-Darby canine kidney (MDCK) cells. Each cell type was seeded in 6-well plates at a density of 5×10^5^ cells per well and incubated at 37 °C for 24 h until approximately 90% confluency was reached. Cells were then inoculated with rTX2/24, rTX2/24-NLuc, rTX2/24-miniGFP2 or rTX2/24-UnaG at a target dose of 30 plaque-forming units (PFU) per well, based on PFU titers determined in Cal-1 cells. Virus adsorption was performed for 1 h at 37 °C. Following adsorption, the inoculum was removed and each well was overlaid with 2 mL of semi-solid medium consisting of 2X complete growth medium mixed with 1% SeaKem agarose, resulting in final concentrations of 1X medium and 0.5% agarose. After polymerization of the overlay, plates were incubated at 37 °C for 72 h. The agarose overlay was then carefully removed, and cells were fixed with 3.7% formaldehyde for 30 min and stained with 0.5% crystal violet solution for 10 min at room temperature. Plaques were visualized and quantified using ViralPlaque software [29].

### Pathogenesis study in mice

Eight-week-old BALB/c mice (*n* = 25) were purchased from The Jackson Laboratory (USA) and housed in HEPA-filtered isolators (Tecniplast BCU) with individual ventilation systems in the animal biosafety level 3 (ABSL-3) facility at the East Campus Research Facility (ECRF), Cornell University. On day 0, mice (*n* = 5 per group) were anesthetized and inoculated intranasally with 50 µL of virus suspension containing 1×10^3^ plaque-forming units (PFU) of one of the following recombinant viruses: rTX2/24, rTX2/24-NLuc, rTX2/24-miniGFP2 or rTX2/24-UnaG. Mock-infected control mice (*n* = 5) were inoculated with cell culture medium. Body weight, clinical signs, morbidity and mortality were monitored and recorded daily. The clinical severity of the inoculated animals was assessed based on a standardized scoring system described previously [30]. Clinical signs such as hunched posture (3 points), ruffled fur (piloerection) (3 points), loss of appetite (2 points), weight loss greater than 20% (10 points) and neurological symptoms (hind-limb paralysis) (10 points) were considered in the scoring system. Daily clinical scores were recorded individually and summed for each group and expressed as average daily group clinical scores. Oropharyngeal swabs were collected using sterile rayon-tipped swabs on days 0, 1, 3, and 5 post-infection (p.i.), placed in 1 mL of virus transport medium (VTM; Corning®, Glendale, AZ, USA), and stored at −80 °C until further analysis. Animals reaching humane endpoints, defined as ≥20% loss of initial body weight, were euthanized and subjected to necropsy and tissue collection. Tissues including brain, nasal turbinate, trachea, lungs, heart, liver, spleen, kidney and intestines were aseptically collected for virus quantification, placed in sterile containers, and stored at −80 °C until processing. Representative sections of each tissue were also fixed in 10% neutral-buffered formalin and processed for histopathological examination. All animal study procedures were reviewed and approved by the Institutional Animal Care and Use Committee of Cornell University (IACUC approval number 2024-0094).

### RNA extraction and RT-PCR

Viral RNA was extracted from oropharyngeal swab samples and 10% (w/v) tissue homogenates using the IndiMag Pathogen Kit (INDICAL Bioscience) on the IndiMag 48s automated nucleic acid extractor (INDICAL Bioscience, Leipzig, Germany), according to the manufacturer’s instructions. Real-time reverse transcription PCR (rRT-PCR) was performed using the Path-ID Multiplex One-Step RT-PCR Kit (ThermoFisher Scientific, Waltham, MA, USA) with primers and probes targeting the influenza A virus matrix (M) gene. Thermal cycling conditions were as follows: reverse transcription at 48 °C for 15 min, initial denaturation at 95 °C for 10 min, followed by 40 cycles of denaturation at 95 °C for 15 s and annealing/extension at 60 °C for 60 s. A standard curve was generated using RNA extracted from HPAI H5N1 TX2/24 virus-spiked medium. Serial 10-fold dilutions of the virus stock were prepared in DMEM, subjected to RNA extraction, and analyzed by rRT-PCR as described above. Cycle threshold (Ct) values were used to estimate viral RNA copy numbers in test samples using relative quantification.

### Virus titrations

All oropharyngeal swab samples and tissue homogenates were subjected to endpoint virus titration using bovine uterine epithelial (Cal-1) cells. Serial 10-fold dilutions of samples were prepared in minimum essential medium (MEM) and inoculated onto Cal-1 cells seeded in 96-well plates. At 48 h post-inoculation, culture supernatants were removed and cells were fixed with 3.7% formaldehyde and processed for immunofluorescence assay (IFA) using an anti-influenza A nucleoprotein (NP) mouse monoclonal antibody (HB65), followed by incubation with an Alexa Fluor 594-conjugated anti-mouse secondary antibody [27, 28]. Virus titers at each time point were determined by endpoint dilution assays and calculated using the Spearman-Kärber method, and titers were expressed as TCID_50_.mL^-1^.

### Virus neutralization assay

Neutralizing antibodies against HPAI H5N1 in serum of uninfected and naturally infected cows was assessed by virus neutralization (VN) assay using the two fluorescent viruses rTX2/24-miniGFP2 and rTX2/24-UnaG and compared with the parental rTX2/24 virus as previously described [26]. For this, serial two-fold serum dilutions (1:8 to 1:1,024) of each serum sample was prepared in MEM and were incubated with 200 TCID_50_ of either rTX2/24, rTX2/24-miniGFP2 or rTX2/24-UnaG virus for 1 h at 37 °C. Next, 100 µL of a cell suspension of Cal-1 cells was added to each well of a 96-well plate and incubated at 37°C for 48 h. Cells inoculated with fluorescent viruses were visualized using a fluorescence microscope (Hybrid microscope ECHO Revolve 3K) to determine neutralizing antibody (NA) titers. While cells inoculated with rTX2/24 were fixed in 3.7% formaldehyde solution after 48 hours of incubation and processed for IFA using anti-influenza A nucleoprotein (NP) mouse monoclonal antibody (HB65), followed by incubation with an Alexa Fluor 594-conjugated anti-mouse secondary antibody. The neutralizing antibody titers were expressed as the reciprocal of the highest serum dilution capable of completely inhibiting HPAI H5N1 virus replication based on the expression of reporter genes or NP positive cells.

### Antiviral assay

The potential application of the H5N1 rTX2/24-NLuc reporter virus for antiviral screens was evaluated against oseltamivir (neuraminidase inhibitor) *in vitro*. Bovine uterine epithelial cells Cal-1 seeded in 96-well plates were infected with rTX2/24-NLuc at a multiplicity of infection (MOI) of 0.01. Virus adsorption was carried out at 37 °C for 1 h, after which the inoculum was removed and replaced with 0.05 mL of MEM supplemented with 2% fetal bovine serum containing oseltamivir at concentrations ranging from 0.009 to 20 mM. Plates were then incubated at 37 °C. Infected but untreated cells served as virus controls. At 24 h post-infection, culture supernatants were removed, and cells were lysed with 1X Passive Lysis Buffer (Promega, USA) for 10 min. Luminescence was quantified using Nano-Glo luciferase assay kit (Promega, USA). The percent inhibition value was calculated to obtain the 50% inhibitory concentration (IC_50_).

### *In situ* RNA detection

Formalin-fixed, paraffin-embedded tissue samples were sectioned at 5 µm and analyzed by *in situ* hybridization (ISH) using RNAscope® ZZ probe technology (Advanced Cell Diagnostics, Newark, CA, USA). Tissues from virus-inoculated and mock-infected control mice were processed using the RNAscope® 2.5 HD Reagents-RED kit (Advanced Cell Diagnostics) in combination with the V-InfluenzaA-H5N8-M2M1 probe (Advanced Cell Diagnostics), which targets influenza A H5Nx clade 2.3.4.4b viruses, according to the manufacturer’s instructions. ISH signals were amplified using a series of alkaline phosphatase-conjugated amplifiers, visualized by incubation with a red chromogenic substrate at room temperature for 10 min, and counterstained with hematoxylin.

### Accession number

All sequence data from the reporter viruses generated in this study were submitted to the NCBI BioProject (BioProject ID PRJNA1406649).

### Statistical analysis and data plotting

Statistical analysis was performed by One-way or 2-way analysis of variance (ANOVA) followed by multiple comparisons. Correlation coefficient (R^2^) values were calculated to compare the sensitivity of the recombinant viruses in detecting neutralizing antibody titers in the serum. Statistical analysis and data plotting were performed using the GraphPad Prism software (version 9.0.1).

## Results

### Generation of recombinant HPAI H5N1 viruses expressing NLuc, miniGFP2 and UnaG reporters

A reverse genetics system for the bovine H5N1 virus based on an isolate A/Cattle/Texas/06322424-1/2024 (TX2/24) obtained from milk from infected dairy cows was established in our laboratory and used as backbone to generate a recombinant virus expressing the nanoluciferase (rTX2/24-NLuc), miniGFP2 (rTX2/24-miniGFP2) and UnaG (rTX2/24-UnaG) reporter proteins. The NS segment of the TX2/24 virus was modified to encode NLuc, miniGFP2 or UnaG fusion protein from a single non-overlapping transcript. The reporter genes were cloned at the C-terminal of NS1 and the NS1 and NEP open reading frames were separated by the porcine teschovirus 1 2A autoproteolytic cleavage site (**Fig. 1A**). Full length genome sequences and modified NS segment of the TX2/24 virus were synthesized commercially and cloned into dual promoter pHW2000 vector (**Fig. 1B**). Recombinant viruses were rescued upon co-transfection of seven TX2/24 gene segments (PB2, PB1, PA, HA, NA, NP and M) and the modified NS-NLuc, NS-miniGFP2 or NS-UnaG segment into a co-culture of HEK293T and Cal-1 cells (**Fig. 1C**). Working stocks of the recombinant viruses were prepared in 9-10-day old embryonated chicken eggs. Viruses from the initial rescue and working stocks were sequenced and no unwanted mutation was found. Presence of infectious viruses (rTX2/24-miniGFP and rTX2/24-UnaG) and expression of fluorescence proteins were confirmed upon inoculation of Cal-1 cells and immunofluorescence imaging, while rescue of the nano luciferase expressing virus was confirmed by immunofluorescence using an NP-specific antibody (**Fig. 1D**). Both rTX2/24-miniGFP2 and rTX2/24-UnaG infected cells showed bright green signal in the infected cells (**Fig. 1D**), while the rTX2/24-NLuc expressed high levels of luciferase as measured by luminescence (**Fig. 1E**).

**Fig. 1.**
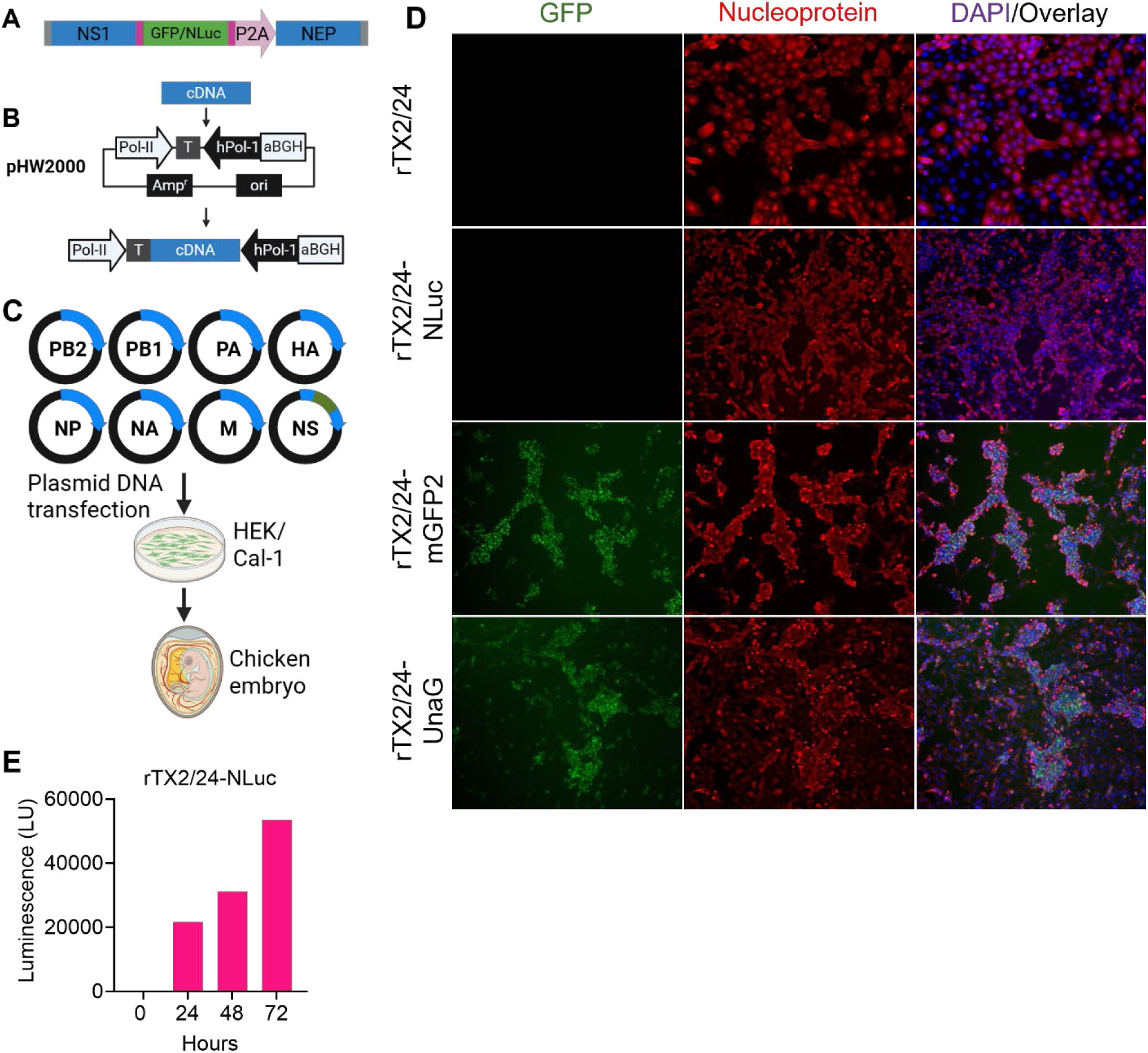
Generation of recombinant H5N1 TX2/24 viruses expressing bioluminescence and fluorescence reporter proteins. (A-C) Schematics of influenza virus reverse genetics system to generate reporter viruses. (D) Immunofluorescence images showing viral nucleoprotein and GFP signals in Cal-1 cells inoculated with recombinant parental and reporter viruses. (E) Luciferase reporter assay on rTX2/24-NLuc-infected Cal-1 cell culture supernatant and expressed as luminescence (LU), n = 1.

### Recombinant H5N1 TX2/24 bioluminescent and fluorescent reporter viruses exhibited efficient replication but reduced cell-to-cell spread *in vitro*

To evaluate virus replication kinetics, we performed multicycle growth curve analyses of the three reporter viruses in bovine uterine epithelial cells (Cal-1), human lung adenocarcinoma cells (A549), and Madin-Darby canine kidney (MDCK) cells, and compared them with the parental rTX2/24 virus. All three reporter viruses replicated efficiently in all tested cell lines, showing comparable titers to the parental rTX2/24 virus (**Fig. 2A**). To quantify viral loads, we performed a luciferase reporter assay on samples from growth curve experiments shown in **Fig. 2A**. The luciferase assay revealed growth kinetics for rTX2/24-NLuc that closely mirrored those obtained using the infectious virus titration method (**Fig. 2B**).

**Fig. 2.**
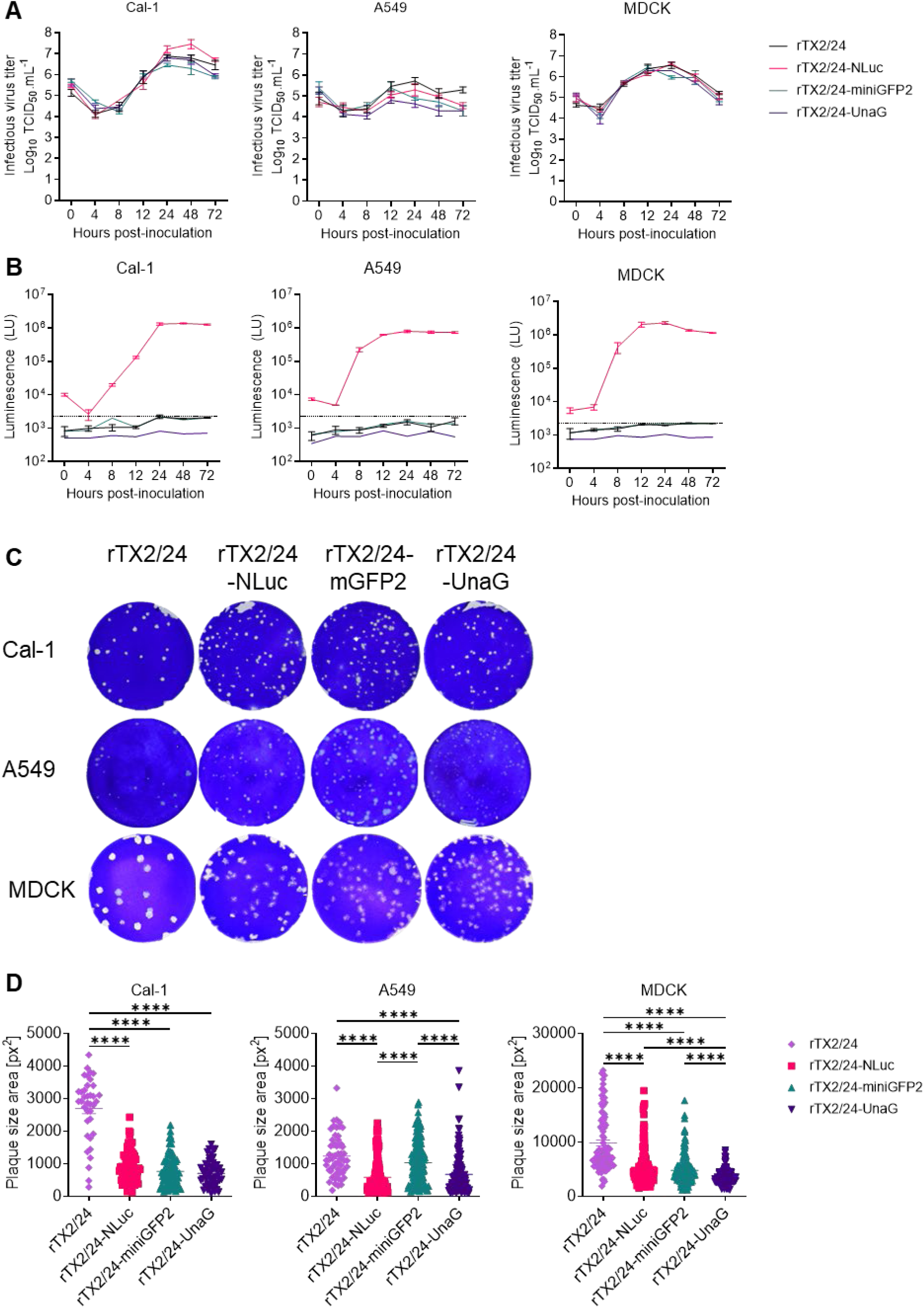
Replication kinetics and plaque size phenotype of HPAI H5N1 reporter viruses in mammalian cells. (A) Multicycle growth curves. Bovine uterine epithelial cells (Cal-1), human lung adenocarcinoma (A549) cells and Madin-Darby canine kidney (MDCK) cells were infected (MOI 0.1) with HPAI H5N1 rTX2/24, rTX2/24-NLuc, rTX2/24-miniGFP2 and rTX2/24-UnaG viruses and virus titers were determined at indicated time points by limiting dilution method and expressed as TCID_50_.mL^-1^. The limit of detection was 10^1.05^ TCID_50_.mL^-1^ (B) Luciferase reporter assays on growth curve samples from (A) and expressed as luminescence (LU). Dashed lines indicate background luminescence. (A-B) Data indicates mean ± SEM, n = 3, three independent experiments. (C) Viral plaque phenotype. Cal-1, A549 and MDCK cells were infected (30 plaque forming unit/well) with recombinant H5N1 viruses and overlaid with medium containing agar. Plates were incubated at 37 °C for 72 h, the agar overlay was removed, cells were fixed, and the monolayer was stained with 0.5% crystal violet. (D) The diameters of viral plaques from Figure C were measured using ViralPlaque software in pixel^2^. One-way ANOVA with Tukey’s multiple comparison test, **** *p* < 0.0001.

We next examined the plaque morphologies of the recombinant viruses in Cal-1, A549, and MDCK cells. In most cases, all three reporter viruses produced smaller plaques than the parental rTX2/24 virus, indicating reduced cell-to-cell spread (**Fig. 2C-2D**). Among the three reporter viruses, the UnaG-expressing virus displayed the greatest attenuation in cell-to-cell spread, while rTX2/24-NLuc and rTX2-miniGFP2 produced comparable plaque sizes in both Cal-1 and MDCK cells (**Fig. 2C-2D**). Collectively, these results demonstrate that all three recombinant reporter viruses are replication-competent but exhibit reduced cell-to-cell spread relative to the parental rTX2/24 virus *in vitro*.

### Recombinant H5N1 TX2/24 reporter viruses caused lethal infection in mice but showed reduced virus shedding

To assess the pathogenicity of the reporter viruses, mice were inoculated intranasally and monitored for clinical signs, body temperature, and virus shedding in oropharyngeal swabs (**Fig. 3A**). Clinical severity was evaluated using a standardized scoring system [30] and compared among mice inoculated with the three reporter viruses and the parental rTX2/24 virus. Mice infected with either parental or reporter viruses developed typical clinical signs of HPAI H5N1 infection, including hunched posture, ruffled fur, and inappetence beginning at day 2 post-inoculation (pi), which progressed to more severe disease (**Fig. 3B**). The clinical sign index (CSI) values were comparable across all infected groups, except for the rTX2/24-UnaG group, which exhibited slightly lower scores on days 2 and 3 pi. While mock-inoculated mice maintained stable body weight, all virus-inoculated mice experienced rapid weight loss, reaching the humane endpoint (≥20% body weight loss) by day 4 pi, except for one mouse in the rTX2/24-miniGFP2 group that reached the endpoint on day 6 pi (**Fig. 3C**). Subcutaneous implantation of biocompatible temperature transponders allowed continuous monitoring of body temperature changes. Although some infected mice exhibited transient fever (up to +1.7°C), most showed a rapid decline in body temperature concurrent with disease progression and increased clinical disease severity (**Fig. 3D**). Ultimately, all infected animals reached humane endpoints by day 4 pi, except for one rTX2/24-miniGFP2-inoculated mouse, which succumbed on day 6 pi (**Fig. 3E**).

**Fig. 3.**
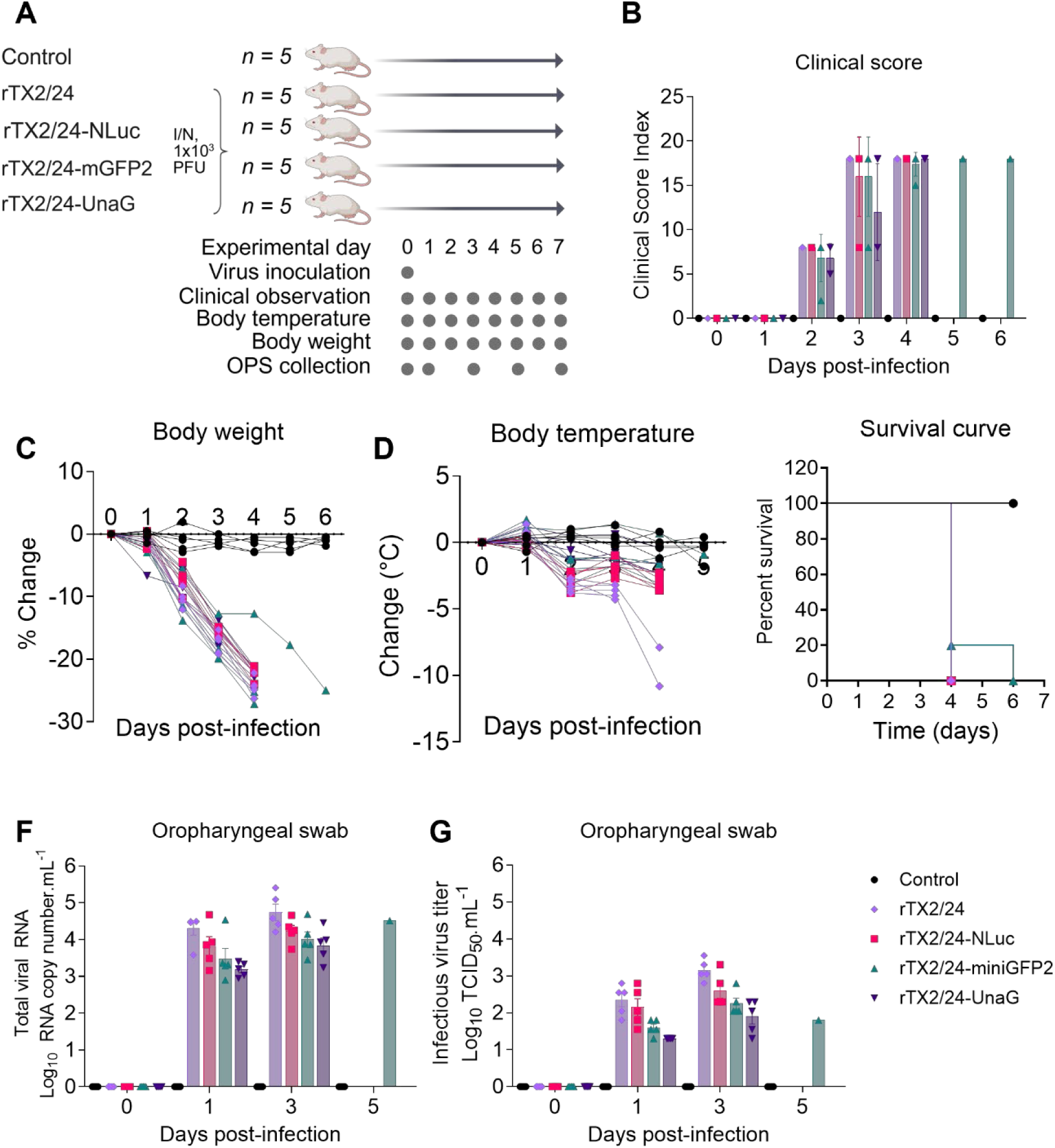
Pathogenesis of recombinant HPAI H5N1 viruses in mice. (A) Experimental design and monitoring and sampling schedule. BALB/c mice (*n* = 25) were mock inoculated (*n* = 5) or inoculated intranasally with 1×10^3^ PFU of HPAI H5N1 rTX2/24 (*n* = 5), rTX2/24-NLuc (*n* = 5), rTX2/24-miniGFP2 (*n* = 5) and rTX2/24-UnaG (*n* = 5) viruses and changes in clinical severity (B) body weight gain (C) and body temperature (D) were monitored for 6 days post-infection (pi). (E) Survival curve. HPAI H5N1 viral RNA (F) and infectious viral (G) loads quantified by rRT-PCR and virus titration in oral secretions collected on days 0, 1, 3 and 5 pi. Virus titers were determined using endpoint dilutions and expressed as TCID_50_.mL^-1^. The limit of detection (LOD) for infectious virus titration was 10^1.05^ TCID_50_.mL^-1^.

To assess viral shedding, HPAI H5N1 RNA and infectious virus titers were quantified in oropharyngeal swabs by rRT-PCR and virus titrations, respectively. The parental rTX2/24 and bioluminescent rTX2/24-NLuc viruses exhibited comparable viral RNA, and infectious virus loads throughout the experimental period, whereas both fluorescent reporter viruses (rTX2/24-miniGFP2 and rTX2/24-UnaG) showed reduced shedding (**Fig. 3F-3G**). Notably, the two fluorescent viruses demonstrated similar shedding patterns.

### Tropism and tissue distribution of HPAI H5N1 virus in mice

To characterize the tissue tropism and distribution of recombinant HPAI H5N1 viruses, mice were inoculated with the parental rTX2/24 virus or one of three reporter variants (rTX2/24-NLuc, rTX2/24-miniGFP2, and rTX2/24-UnaG). Viral loads were quantified by real-time reverse transcription PCR (rRT-PCR), virus titration, and *in situ* hybridization across multiple organs, including the brain, nasal turbinate, trachea, lung, heart, liver, spleen, kidney, and small intestine. Overall, both viral RNA and infectious virus titers were comparable among the parental and reporter viruses in respiratory tissues (**Fig. 4A-B**). The highest viral RNA concentrations were detected in the lungs (7.93-8.47 log_10_ genome copies.g^-1^), followed by nasal turbinates (6.68-7.13 log_10_ genome copies.g^-1^) and trachea (6.09-6.6 log_10_ genome copies.g^-1^) (**Fig. 4A**). Notably, the heart exhibited approximately 2-log_10_ higher viral RNA levels in rTX2/24-inoculated mice (7.34 ± 0.40 log_10_ genome copies.g^-1^) than in reporter virus-inoculated mice, with the lowest RNA loads observed in the rTX2/24-UnaG group (4.75 ± 0.46 log_10_ genome copies.g^-1^). Moderate viral RNA levels were detected in the brain, which were significantly higher (p ≤ 0.05) in parental rTX2/24-inoculated mice (7.33 ± 0.63 log_10_ genome copies.g^-1^) compared with the reporter virus groups (5.53-6.45 log_10_ genome copies.g^-1^). Relatively higher amounts of viral RNA were also detected in the liver, spleen, and kidney of rTX2/24-inoculated mice, and these levels were significantly higher (p ≤ 0.05) than those detected in reporter virus-infected animals.

**Fig 4.**
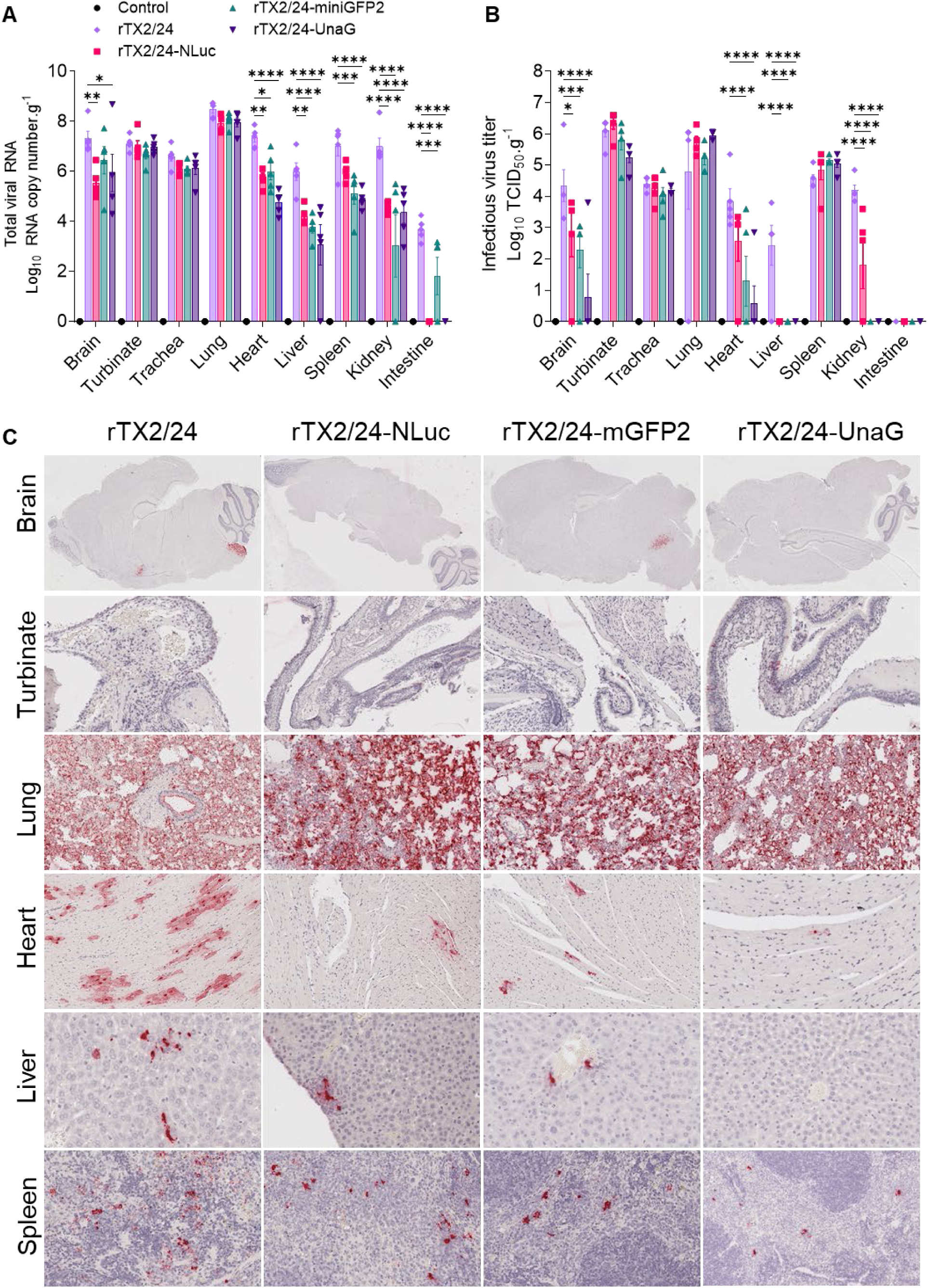
Virus distribution in tissues of mice infected with recombinant HPAI H5N1 viruses. HPAI H5N1 viral RNA (A) and infectious virus (B) loads quantified by rRT-PCR and virus titration in tissues of mice collected at necropsy. Virus titers were determined using endpoint dilutions and expressed as TCID_50_.g^-1^. *In situ* hybridization (ISH) labeling (C, red color) of viral RNA in tissues of HPAI H5N1 infected mice. Representative images of 5 animals per group were shown. (A-B) Data indicate mean ± SEM of 5 mice per group per tissues. Two-way ANOVA with Tukey’s multiple comparison test, * *p* < 0.05, ** *p* < 0.01, *** *p* < 0.001, and **** *p* < 0.0001.

Quantification of infectious virus titers corroborated the rRT-PCR findings (**Fig. 4B**). High infectious virus titers were observed in the nasal turbinates (5.25-6.35 log_10_ TCID_50_.g^-1^) and lungs (4.79-5.95 log_10_ TCID_50_.g^-1^), with no significant differences among the groups. Moderate titers were detected in the trachea (4.05-4.40 log_10_ TCID_50_.g^-1^), which was also comparable between viruses. Among non-respiratory tissues-including the brain, heart, liver, spleen, and kidney-moderate levels of infectious virus were recovered from mice inoculated with both rTX2/24 and rTX2/24-NLuc, whereas only low to undetectable levels were found in mice inoculated with the fluorescent reporter viruses rTX2/24-miniGFP2 and rTX2/24-UnaG (**Fig. 4B**). No infectious virus was detected in the small intestine of any inoculated animal.

*In situ* hybridization revealed intense hybridization signals in the lungs across all four inoculated groups (**Fig. 4C**). Hybridization signals were distributed throughout the lung parenchyma, including the alveolar epithelium and lumen, endothelium, and bronchial epithelial cells. Moderate hybridization signals were also observed in the nasal turbinates and trachea. A higher number of myocardial cells exhibited hybridization signals in rTX2/24-inoculated mice, whereas only a few positive cells were observed in the reporter virus groups. Viral RNA labeling in the spleen and brain was low but consistently higher in rTX2/24-inoculated mice compared with those infected with reporter viruses. In the brain, focal labeling was detected in the cerebrum and localized to neurons, astrocytes, and endothelial cells.

Collectively, these results indicate that the H5N1 rTX2/24 virus replicates efficiently in the respiratory tract and is capable of systemic dissemination, particularly to the brain and heart. In contrast, the reporter viruses, especially the fluorescent reporters, exhibit reduced replication in extrapulmonary tissues, suggesting that reporter gene insertion may attenuate systemic spread without substantially affecting replication in the respiratory tract.

### Application of reporter viruses for sensitive quantification of serum neutralizing antibodies and antiviral screening

We applied these two fluorescent reporter viruses in virus neutralization assays to measure neutralizing antibody responses in a panel of 30 HPAI H5N1-positive and 15 negative cattle serum samples. Neutralization assays were performed using recombinant viruses expressing miniGFP2 (rTX2/24-miniGFP2) or UnaG (rTX2/24-UnaG) and the results were compared with those obtained using the parental rTX2/24 virus. Both fluorescent reporter viruses accurately discriminated between positive and negative sera, and the neutralizing antibody titers were comparable across all viruses, with strong correlations (R^2^ = 0.9376-0.9727) (**Fig. 5A-C**). We further evaluated the potential of the nanoluciferase reporter virus rTX2/24-NLuc for antiviral screening against oseltamivir (neuraminidase inhibitor). Oseltamivir showed a dose-dependent inhibition in the luminescence signals in rTX2/24-NLuc virus-infected cells with a 50% inhibitory concentration (IC_50_) of 295 µM. Collectively, these results demonstrate that fluorescent reporter viruses are reliable tools for serological diagnostics and are well suited for high-throughput neutralization and antiviral assays.

**Fig 5.**
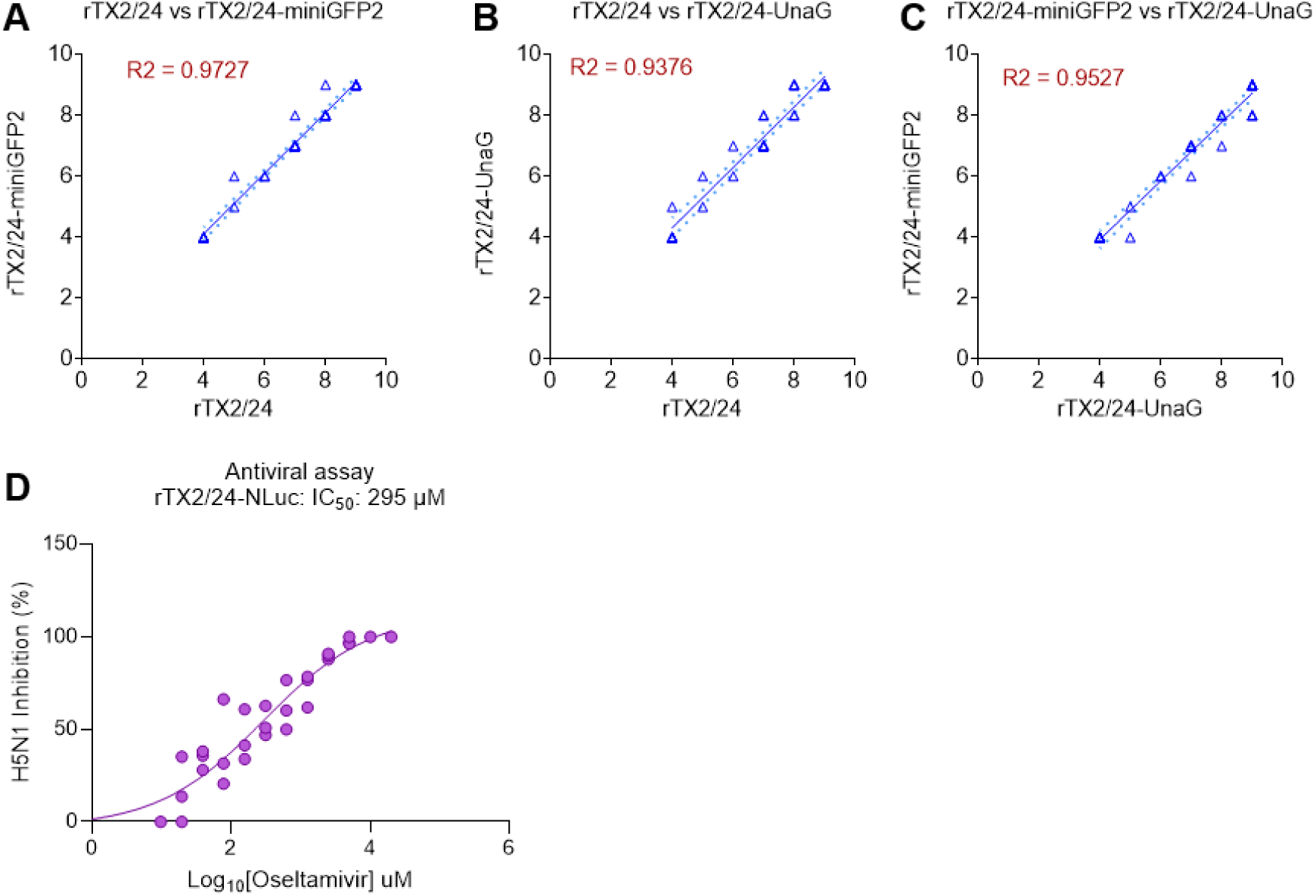
Evaluation of fluorescent reporter viruses in virus neutralization and antiviral assays. (A-C) Serum samples from HPAI H5N1 infected (n = 30) and control (n = 15) farms were tested for the presence of H5N1-specific neutralizing antibodies using virus neutralization assays. Neutralizing antibody titers obtained with recombinant fluorescent reporter viruses expressing miniGFP2 (rTX2/24-miniGFP2) or UnaG (rTX2/24-UnaG) were compared with those obtained using the parental rTX2/24 virus. (D) Antiviral efficacy of oseltamivir against rTX2/24-NLuc virus. Cal-1 cells were infected with rTX2/24-NLuc virus using MOI 0.01 followed by treatment with increasing doses of oseltamivir (0.009–20 mM) for 24 h. Cells were lysed and luminescence was measured. The percent inhibition value was calculated to obtain the 50% inhibitory concentration (IC_50_). n = 3.

## Discussion

The continued emergence of clade 2.3.4.4b H5N1 viruses in mammalian hosts, including dairy cattle and humans underscores the need for biologically relevant reporter systems that enable quantitative assessment of viral replication, pathogenesis and transmission. In this study, we generated three replication-competent reporter H5N1 viruses expressing nanoluciferase (NLuc), miniGFP2, or UnaG using a bovine-derived TX2/24 backbone representative of currently circulating B3.13 genotype viruses. Our results demonstrate that these reporter viruses retain robust capability replication *in vitro* and *in vivo* while providing sensitive and versatile readouts suitable for antiviral and serological applications.

Luciferase reporter viruses offer distinct advantages for *in vivo* studies, particularly for non-invasive and longitudinal monitoring, compared with GFP-based reporter viruses [12, 13, 17, 23, 31]. In contrast, reporter viruses containing full-length GFP sequences (∼ 0.7 kb) substantially increase the length of the NS segment, often resulting in viral attenuation and loss of reporter stability following serial passage in cell culture or *in vivo* [17, 32]. To overcome these limitations, we developed two fluorescent reporter viruses expressing miniGFP2, a 13 kDa green fluorescent protein with improved photochemical stability that enables long-term live-cell imaging [24], and UnaG, a 15.6 kDa bilirubin-inducible fluorescent protein [25].

The reporter genes were inserted into within the NS segment open reading frame to generate NS1-reporter-2A-NEP constructs, allowing expression of NS1 fused to the reporter protein and independent expression of a functional NEP following autoproteolytic cleavage at the 2A site [32]. Preservation of NEP function is critical for efficient viral replication and nuclear export of viral ribonucleoproteins. This strategy successfully maintained multicycle replication competence of the recombinant viruses in bovine, human and canine cells, consistent with previous studies using similar designs [12, 13, 31–33]. Moreover, the strong correlation between NLuc luminescence and infectious virus titers validates rTX2/24-NLuc as a reliable surrogate for conventional virus quantification and supports its utility in rapid, high-throughput antiviral screening platforms.

Despite comparable replication kinetics, all reporter viruses exhibited reduced plaque sizes relative to the parental rTX2/24 virus, indicative of impaired cell-to-cell spread *in vitro*. This phenotype was most pronounced for the UnaG-expressing virus, likely reflecting modest fitness costs associated with increased NS segment length, altered NS1 expression levels, or incomplete 2A cleavage efficiency [34]. Although previous studies have shown that NS1-GFP fusion viruses can effectively inhibit interferon-β induction to levels comparable to wild-type virus [32], similar reductions in plaque size have been reported for other influenza reporter viruses and generally do not preclude their utility in pathogenesis or immunological studies when overall replication competence is preserved [14], a feature that is observed in the reporter viruses developed here.

In the mouse model, all reporter viruses caused severe, lethal disease with clinical progression comparable to that of the parental virus, demonstrating that reporter insertion did not abolish virulence in this highly susceptible host. However, the fluorescent reporter viruses displayed reduced viral shedding and modest attenuation of systemic dissemination compared with rTX2/24 and rTX2/24-NLuc, consistent with previous observations [14, 32]. While viral replication in the respiratory tract was largely preserved, extrapulmonary spread, particularly to the brain and heart, was reduced in mice infected with the fluorescent reporter viruses. These findings suggest that reporter gene insertion preferentially impacts viral fitness in systemic tissues without substantially altering replication at the primary site of infection. Given the critical role of NS1 in antagonizing host innate immune responses, subtle changes in NS1 function or expression may disproportionately affect viral spread in the more restrictive immune environments of extrapulmonary tissues [35]. However, the spatial and temporal dynamics of virus replication of the reporter viruses in tissues should be further assessed using the *in vivo* imaging system (IVIS).

A key advantage of the reporter viruses described here is their application to neutralization assays using sera from naturally infected dairy cattle and screening antivirals using high throughput assays. Both miniGFP2- and UnaG-expressing viruses accurately quantified neutralizing antibody titers and showed strong concordance with assays performed using the parental virus. These results demonstrate that fluorescent reporter viruses can serve as reliable surrogates for wild-type H5N1 in serological assays, enabling higher throughput and reduced assay time without compromising accuracy. Such tools will be particularly valuable for large-scale surveillance, vaccine evaluation, and antiviral screening efforts in livestock populations affected by H5N1 outbreaks [26, 33].

In conclusion, we developed three replication-competent H5N1 reporter viruses based on a bovine-derived clade 2.3.4.4b isolate (genotype B3.13) that retain key biological properties of the parental virus while providing sensitive and versatile readouts of infection. These reporter systems expand the experimental toolkit for studying contemporary H5N1 viruses and will facilitate mechanistic investigations of viral replication, host adaptation and immune responses in mammalian hosts.

## Data availability statement

All data generated in this study are presented in the article.

## Biosafety and biosecurity, and ethical regulations

All work involving rescuing, handling and propagation of HPAI H5N1 viruses was performed following strict biosafety measures in the Animal Health Diagnostic Center (AHDC) research BSL-3 suite at the College of Veterinary Medicine, Cornell University. The animal study procedures were reviewed and approved by the Institutional Animal Care and Use Committee at Cornell University (IACUC approval number 2024-0094) and conducted in the ABSL-3 suite at the College of Veterinary Medicine, Cornell University. All relevant ethical regulations were followed.

## Acknowledgements

The work was supported by internal grants from the College of Veterinary Medicine, Cornell University. We acknowledge Russel J. R. Barkley, Department of Microbiology and Immunology and Salman L. Butt, Department of Population Medicine and Diagnostic Sciences, College of Veterinary Medicine, Cornell University for their technical support.

